# How a highly acidic SH3 domain folds in the absence of its charged peptide target

**DOI:** 10.1101/2023.03.21.532811

**Authors:** Valeria Jaramillo-Martinez, Matthew J. Dominguez, Gemma M Bell, Megan E Souness, Anna H. Carhart, M. Adriana Cuibus, Elahe Masoumzadeh, Benjamin J Lantz, Aaron J Adkins, Michael P Latham, K. Aurelia Ball, Elliott J Stollar

## Abstract

Charged residues on the surface of proteins are critical for both protein stability and interactions. However, many proteins contain binding regions with a high net-charge that may destabilize the protein but are useful for binding to oppositely charged targets. We hypothesized that these domains would be marginally stable, as electrostatic repulsion would compete with favorable hydrophobic collapse during folding. Furthermore, by increasing the salt concentration we predict that these protein folds would be stabilized by mimicking some of the favorable electrostatic interactions that take place during target binding. We varied the salt and urea concentrations to probe the contributions of electrostatic and hydrophobic interactions for the folding of the 60-residue yeast SH3 domain found in Abp1p. The SH3 domain was significantly stabilized with increased salt concentrations according to the Debye-Huckel limiting law. Molecular dynamics and NMR show that sodium ions interact with all 15 acidic residues but do little to change backbone dynamics or overall structure. Folding kinetics experiments show that the addition of urea or salt primarily affects the folding rate, indicating that almost all the hydrophobic collapse and electrostatic repulsion occurs in the transition state. After the transition state formation, modest yet favorable short-range salt-bridges are formed along with hydrogen bonds, as the native state fully folds. Thus, hydrophobic collapse offsets electrostatic repulsion to ensure this highly charged binding domain can still fold and be ready to bind to its charged peptide targets, a property that is likely evolutionarily conserved over one billion years.

**Statement for broader audience:** Some protein domains are highly charged because they are adapted to bind oppositely charged proteins and nucleic acids. However, it is unknown how these highly charged domains fold as during folding there will be significant repulsion between like-charges. We investigate how one of these highly charged domains folds in the presence of salt, which can screen the charge repulsion and make folding easier, allowing us to understand how folding occurs despite the protein’s high charge.

**Supplementary material:**

- Supplementary material document containing additional details on protein expression methods, thermodynamics and kinetics equations, and the effect of urea on electrostatic interactions, as well as 4 supplemental figures and 4 supplemental data tables. (**Supplementary_Material.docx**), 15 pages
- Supplemental excel file containing covariation data across AbpSH3 orthologs (**FileS1.xlsx**)

## Introduction

Electrostatic charge is a fundamental characteristic of protein structure. Most proteins contain charged residues on their surface that interact with solvent, and many even have patches of net charge that are important for interactions with small molecule ligands, nucleic acid, and other proteins through electrostatic attraction (1). These regions of surface charge often result in proteins with net dipole moments as well as overall net charge. However, this clustering of residues with like charges in the folded state is fundamentally destabilizing due to electrostatic repulsion. Therefore, selection pressure driving clustering of charges on a protein surface competes with the pressure to maintain a stable folded state. Understanding this balance is important for both characterizing protein interaction domains and for engineering new protein interactions. However, attempts to predict how charge affects the stability and structure of proteins has remained a challenge (2), as mutations and changes in pH often have unexpected effects on folding kinetics and thermodynamics.

The role of electrostatic interactions in protein folding has often been probed by varying the salt concentration in the protein environment. For proteins with a high net charge, increasing salt concentration has been shown to stabilize the folded state (3-6). Salt can affect charged domain stability through both direct ion binding and Debye screening, and may also vary according to ion-specific Hofmeister effects (4, 7, 8). Direct ion binding results in a hyperbolic relationship between free energy of folding and salt concentration, whereas the Hofmeister effect leads to a linear relationship with salt concentration (9). Both effects would also depend on the specific salt ion present. Alternatively, Debye screening would result in a linear increase in free energy of unfolding with the square root of the ionic strength of the solution, according to Equation 1:

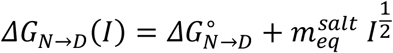

where *I* is the ionic strength of the solution and 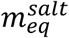 is a constant (9). For highly charged protein domains, salt seems to mainly stabilize through a Debye screening mechanism, which is independent of salt type (3, 5, 9-11).

Electrostatic interactions can be formed either before or after the transition state during the protein folding process and varying the salt concentration can be used to distinguish between these two extremes. In the case of an electrostatically destabilized mutant of thioredoxin, increasing salt concentration stabilized the protein by decreasing the unfolding rate, with little change to the folding rate, indicating that electrostatic interactions occur after transition state formation (6). This contrasts with studies that use charge mutations to enhance protein stability, which appear to primarily operate by increasing the folding rate, while unfolding remains constant (11, 12). For example, charge mutations in the Fyn SH3 domain stabilized the transition state by reducing the repulsive interactions in a nonspecific manner (12). Electrostatic interactions also predominantly occur during transition state formation in a study by Tzul *et al*. that used charge mutations to stabilize four different proteins, leading to more compact, native-like transition states and faster folding (11). Although SH3 domains all share a common fold, the folding and unfolding pathways can vary between different sequences (13). Similarly, electrostatic interactions may play different roles in the folding processes of different SH3 domains, and further study is required to investigate this effect.

In this study, we chose to probe electrostatic interactions in the folding of the SH3 domain from the yeast Actin binding protein 1 (AbpSH3) as it has the highest net negative charge (−12) out of all 28 yeast SH3 domains (14). Furthermore, a comparison with the well-studied SH3 domain from Fyn (net charge of -6) allows us to investigate folding for SH3 domains across a range of net charges. AbpSH3 has been shown to bind to a range of positively charged physiological peptides and peptide mutants, which points to the central role of attractive long-range electrostatics for binding (15-17). Therefore, understanding the folding of AbpSH3 will shed light on the biophysical adaptations necessary for domains that require a high net surface charge for function. SH3 domains are the most common protein-interaction domain in metazoa and are typically composed of 60 residues forming a 5-stranded beta barrel composed of two orthogonal beta sheets packed against each other that bring the N- and C-termini close together. Classically, SH3 domains have been shown to fold via a two-state nucleation condensation process (18-22), whereas others (such as Fyn and PI3K SH3 domains) appear to fold via an intermediate in a multi-state diffusion collision model (4, 23, 24). Recent molecular dynamic simulations have shown there are 2 major types of transition states in SH3 domains (13). In the first unfolding pathway, the dominant secondary structure present in the transition state is between strands 2, 3, and 4. In the second pathway, most secondary structure is between strands 1, 2, and 5. In both pathways, the structured distal loop is present. Currently, it is not known how SH3 domains that conserve very high net charge on their binding surface can remain folded in physiological conditions and how their folding pathway differs from domains with lower net charge.

We first explore the conservation of charged residues in the yeast SH3 domain family, which we recently characterized across a range of species in the fungal kingdom and find that net charge is more conserved than any specific electrostatic interactions that may occur between a pair of residues. We then determine the stability of AbpSH3 at various salt concentrations and show that the domain is destabilized by long-range electrostatic repulsion according to the Debye-Huckel rate limiting law. We show that fast-time scale backbone dynamics in the native-state were not significantly affected, and the overall native state is unaffected by electrostatic repulsion. Importantly, we find that almost all the electrostatic repulsion found in the native state is also found in the transition state and gives insight into AbpSH3 transition state structure and dynamics. Our study sheds light on how a protein interaction domain can successfully fold and accommodate a high net charge that is essential for function.

## Results and Discussion

### The highly charged AbpSH3 is conserved over 1 billion years of evolution

Yeast contains 28 SH3 domains, which we analyzed by sequence alignment across the fungal kingdom. AbpSH3 has the highest net negative surface charge (−12) within the 60-residue domain with 8 aspartates, 7 glutamates, and only 3 lysine residues (**Fig 1a and 3b**). This surface charge, helps AbpSH3 bind to its peptide targets that have high net positive charge such as ArkA, which contains 6 lysine residues. Analysis of our sequence alignment data (14) shows that this high surface charge property is conserved across the fungal kingdom with the average net charge of AbpSH3 orthologs being -10 ± 0.8, resulting in significant electrostatic repulsion (**Fig 1b**). However, the average number of arginine or lysine residues is conserved at 1-1.5, allowing several favorable electrostatic interactions to occur, which appear to be centered around the distal loop that connects strands 3 and 4. For the remaining 27 SH3 domains in yeast, all but one domain conserves some degree of net negative charge (**Fig 1c**). Since most SH3 domains bind to positively charged peptide targets, it is not surprising that a negatively charged functional surface has been conserved. For yeast AbpSH3, the high charge creates a net dipole that facilitates faster association rates (15), which may explain why it has maintained its high surface charge. Overall, yeast AbpSH3 will be destabilized by electrostatic repulsion between acidic residues, although the 3 lysines found away from the binding surface (Fig 3b) appear to offset this by participating in favorable electrostatic interactions (K25-E42, K43-D24, K43-D44, and K47-E40). However, these exact pairwise interactions are not conserved (**File S1**). Significant covariation (using 262 ortholog sequences) only occurs between positions 40 and 47, and for most species, this interaction is between two polar residues and doesn’t require charge-charge interactions. Specific charge-charge interactions are not as important as the net charge and likely the net dipole moment. Although the net number of charged residues is conserved, the only charged residues in yeast AbpSH3 that are conserved, are those on the peptide binding surface. While the charge distribution is clearly important for function, it poses a challenge to the structural stability of the fold, which we aim to explore in this paper.

**Figure 1.**
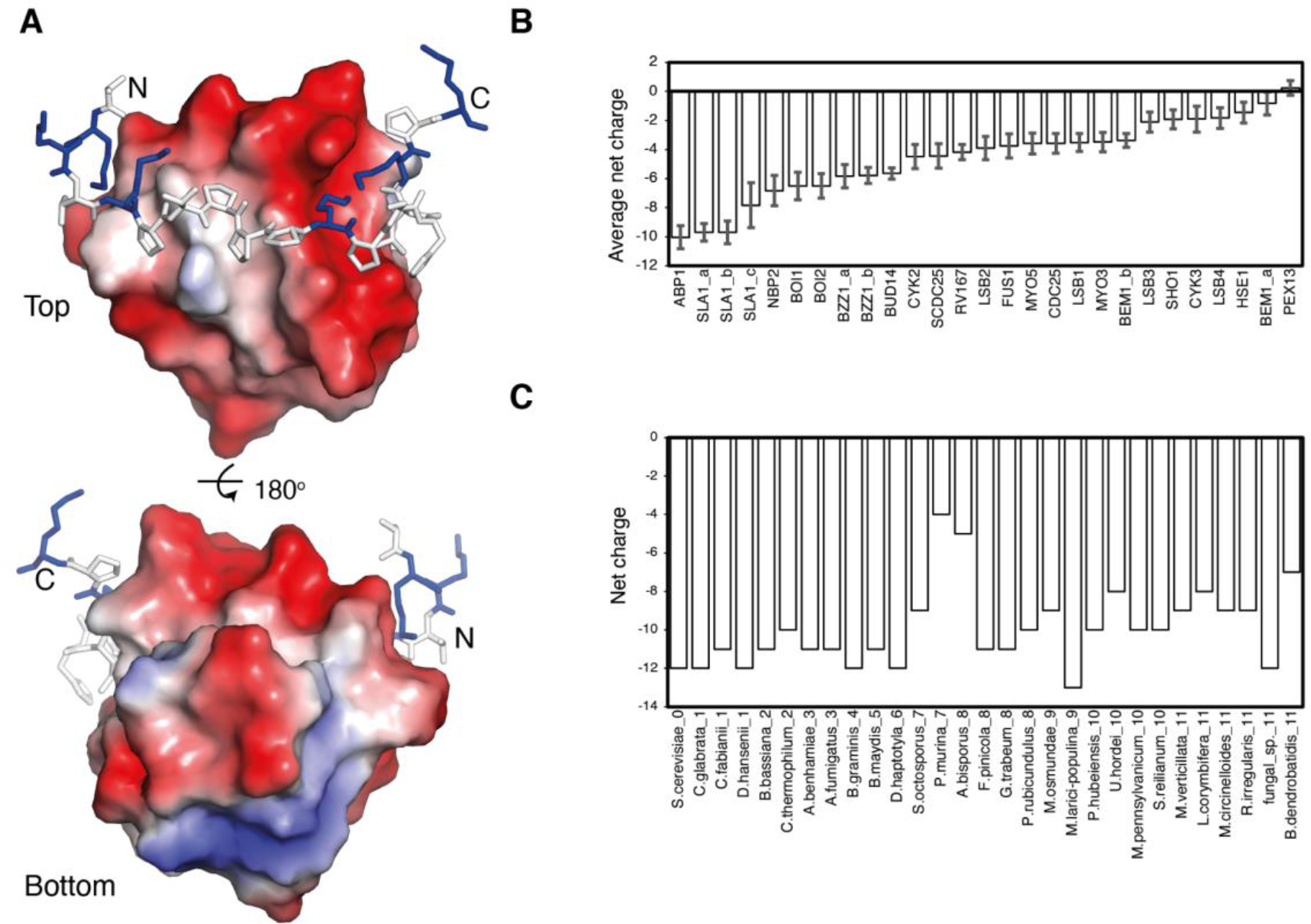
SH3 domains are negatively charged, and this is conserved over evolution. **A**. Electrostatic surface from AbpSH3-ArkA peptide complex (pdb 2rpn). Negative charges are shown in red and positive in blue. **B**. Average net charge across 1 billion years of evolution in the Fungal kingdom for all yeast SH3 domain paralogs. Error bars are 95 % confidence intervals **C**. Average net charge for AbpSH3 orthologs spanning 1 billion years of evolution in the fungal kingdom.

### Long-range electrostatic repulsion during folding destabilizes the domain

The stability of the negatively charged AbpSH3 from *S. cerevisiae* was measured using chemical denaturation and tryptophan fluorescence at a variety of NaCl and KCl concentrations. These salts were chosen as they sit in the middle of the Hofmeister series and are relatively nonchaotropic and nonkosmotropic. This allows us to consider the effect of ionic strength on the stability, while minimizing other confounding effects. We find a significant increase in domain stability (*ΔG*_*N*→*D*_), which is linear with the square root of ionic strength (**Fig 2**). This relationship suggests that the electrostatic repulsion between the 15 acidic residues in AbpSH3 is effectively screened by sodium or potassium cations, according to the Debye-Huckel rate limiting law. We also measured the stability of AbpSH3 from the distant orthologue *B*.*dendrobatidis*, which has a lower net charge of -7 (**Fig 1c**) and also shows a linear relationship with the square root of ionic strength for both salts, evidencing conservation of electrostatic repulsion in AbpSH3 orthologs across the fungal kingdom.

**Figure 2.**
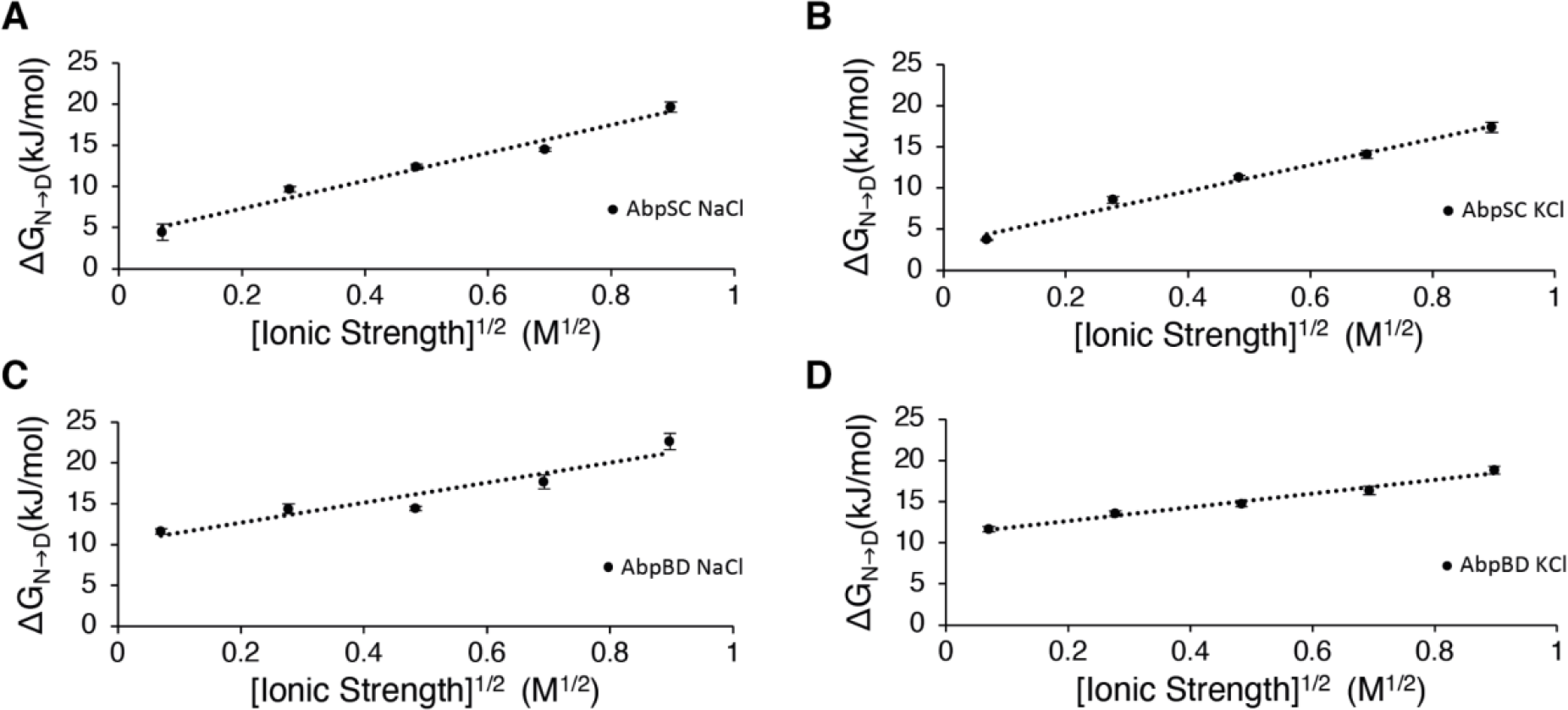
Stability in AbpSH3 from S.cerevisiae (SC) and B.dendrobatidis (BD) increases at higher ionic strength. Free energy of folding shows a linear relationship with the square root of ionic strength. Error bars represent standard deviations across experiment repetitions (some of the error bars are smaller than the markers with values less than ±2 %).

A similar effect for both NaCl and KCl is consistent with Debye-Huckel screening, which is independent of salt type. We do measure a slightly lower stability for AbpSH3 in KCl than in (NaCl 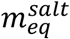is 17.00 for NaCl vs. 15.84 for KCl in AbpSC and 12.24 vs. 8.34 in AbpBD), which may also indicate that direct ion binding or Hofmeister effects slightly contribute to stabilization. However, our domain stability data altogether fits more closely to a linear equation with respect to the square root of ionic strength, than to an equation that is linear with respect to ionic strength or to the log of ionic strength, indicating that the Debye-Huckel equation is a good model for the effect of salt on AbpSH3 folding (**Fig S1 and S2**).

Although the Debye-Huckel limiting law explains how the ionic strength of a salt solution can screen electrostatic interactions non-specifically, MD simulations allow us to observe the specific locations of salt ions in the solution and how they interact with the charged residues in the protein. As expected, positively charged ions spend time within 4 Å of the Asp and Glu charged groups (**Fig 3a**). **Fig 3b** demonstrates that ion interactions are also affected by the net dipole of the SH3 domain, so cations interact more frequently with negatively charged residues near the binding surface at the negative end of the dipole and less frequently with negatively charged residues on the opposite side of the domain, at the more positive end of the dipole. In this way, cations play a similar role to the positively charged partner peptide, ArkA, which is electrostatically attracted to the acidic binding surface. The overall negative charge of the domain also likely contributes to the lower frequency of chloride anion interactions with lysine residues, compared to cation interactions with the negatively charged acidic residues. Potassium cations also directly interact with the acidic SH3 residues although at a lower frequency than sodium. Interestingly, in most cases in the 800 mM potassium chloride simulation there are lower interaction frequencies than the interactions seen in the 300 mM sodium chloride simulations. This is likely due to the increased steric hindrance associated with their larger atomic size, consistent with the thermodynamic data that shows a lower stability of AbpSH3 in KCl than NaCl. Since a typical yeast cell contains ∼300 mM potassium (25), folding will be partially stabilized in these conditions through electrostatic screening, even in the absence of a positively charged binding partner.

**Figure 3.**
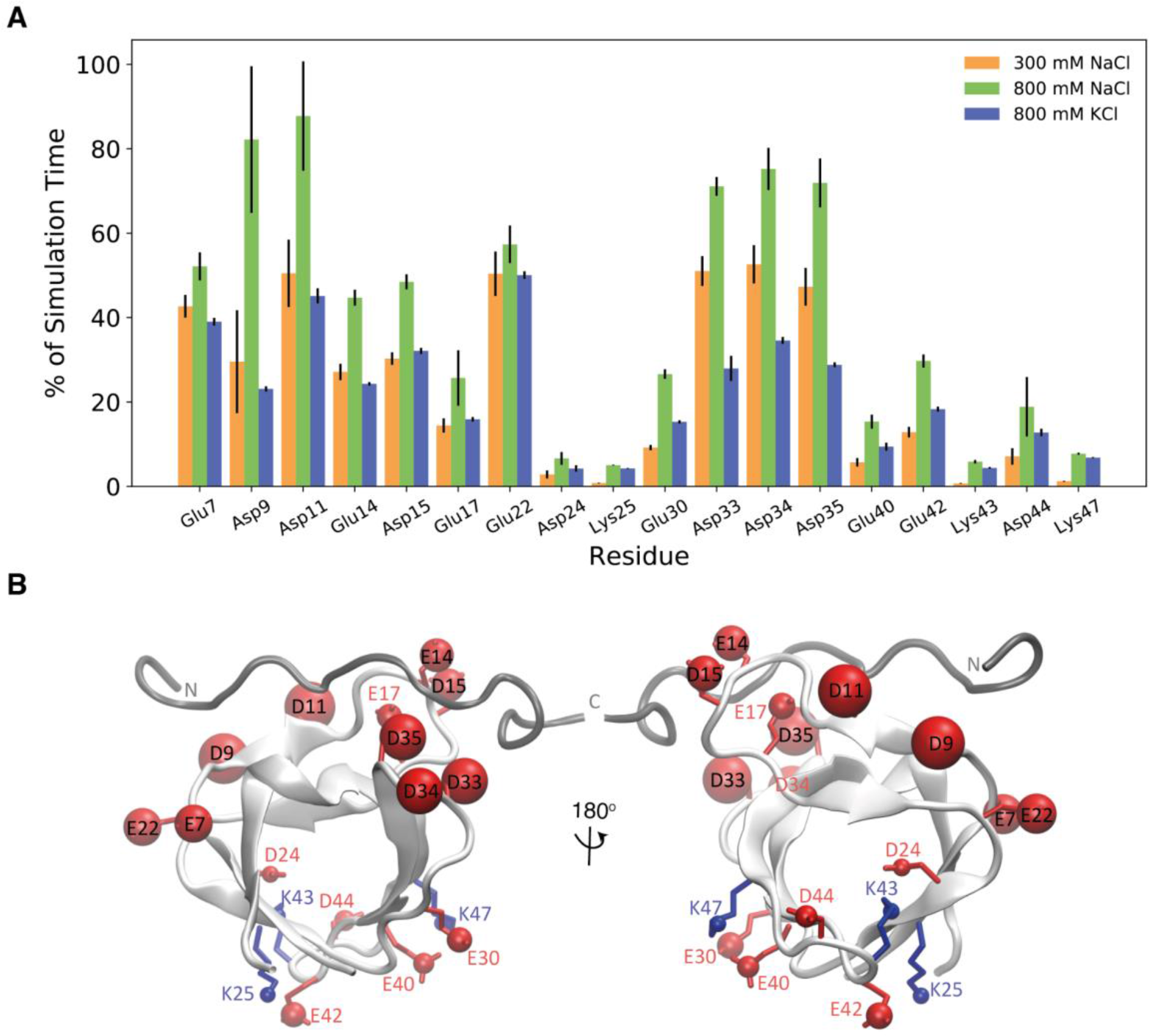
Ion accessibility varies across the domain (in NaCl and KCl) **A**. Percent of simulation time that an ion is in contact with a charged group of a residue in the 300 mM NaCl, 800 mM NaCl, and 800 mM KCl simulations. Interactions with chloride anions are displayed for the positively charged lysine residues and interactions with cations (sodium or potassium) are displayed for the negatively charged aspartate or glutamate residues. **B**. Location of charged residues that interact with ions displayed on AbpSH3-ArkA NMR structure. Positively charged residues are shown in blue and negatively charged residues are shown in red. The ArkA peptide is shown in gray to indicate where it binds, although it was not present in the simulations. The size of the sphere on the charged group of the residue indicates the percent of simulation time that residue is in contact with an oppositely charged ion in the 800 mM NaCl simulations.

### Fast time-scale dynamics and structure were generally unchanged (NMR and MD)

To assess any changes in structure or dynamics that occur due to salt, the NH correlation spectra for AbpSH3 in 0 and 800 mM NaCl were compared. AbpSH3 maintained a well dispersed spectrum in the presence of 800 mM NaCl and small chemical shift perturbations were observed, indicating that salt has a minimal effect on the structure of the native state (**Figure S3**). We also measured ^15^NH R_1_, R_2_, and heteronuclear NOE data to characterize the effects of salt on the pico-to-nanosecond timescale dynamics of AbpSH3. We observed little difference in heteronuclear NOE values, which are sensitive to picosecond timescale motions, upon the addition of salt. Backbone amide order parameters, which report on the amplitude of motion on the fast timescale, were also calculated from the relaxation data, and again, no effect was seen in the presence of salt. Our relaxation data therefore suggests that salt also has little effect on the backbone dynamics of the folded domain.

We also performed MD simulations to examine the fast time-scale dynamics, as well as any conformational changes of the domain under different salt conditions. We performed 10 independent simulations of the AbpSH3 domain under no salt, 800 mM NaCl, and 800 mM KCl conditions. Atomic fluctuations were used to quantify the dynamics of the AbpSH3 backbone during the simulations (**Fig 4c**). Backbone fluctuations were lower in the regions of the sequence involved in β-sheets and higher in loop regions, particularly the RT loop that is part of the peptide binding surface. Overall, we saw no significant difference in backbone fluctuations between the simulations at different salt conditions. We also observed no large changes in the conformation of the SH3 domain in salt compared to no salt, confirming that the folded state of the domain is largely unaffected by the presence of high salt.

**Figure 4.**
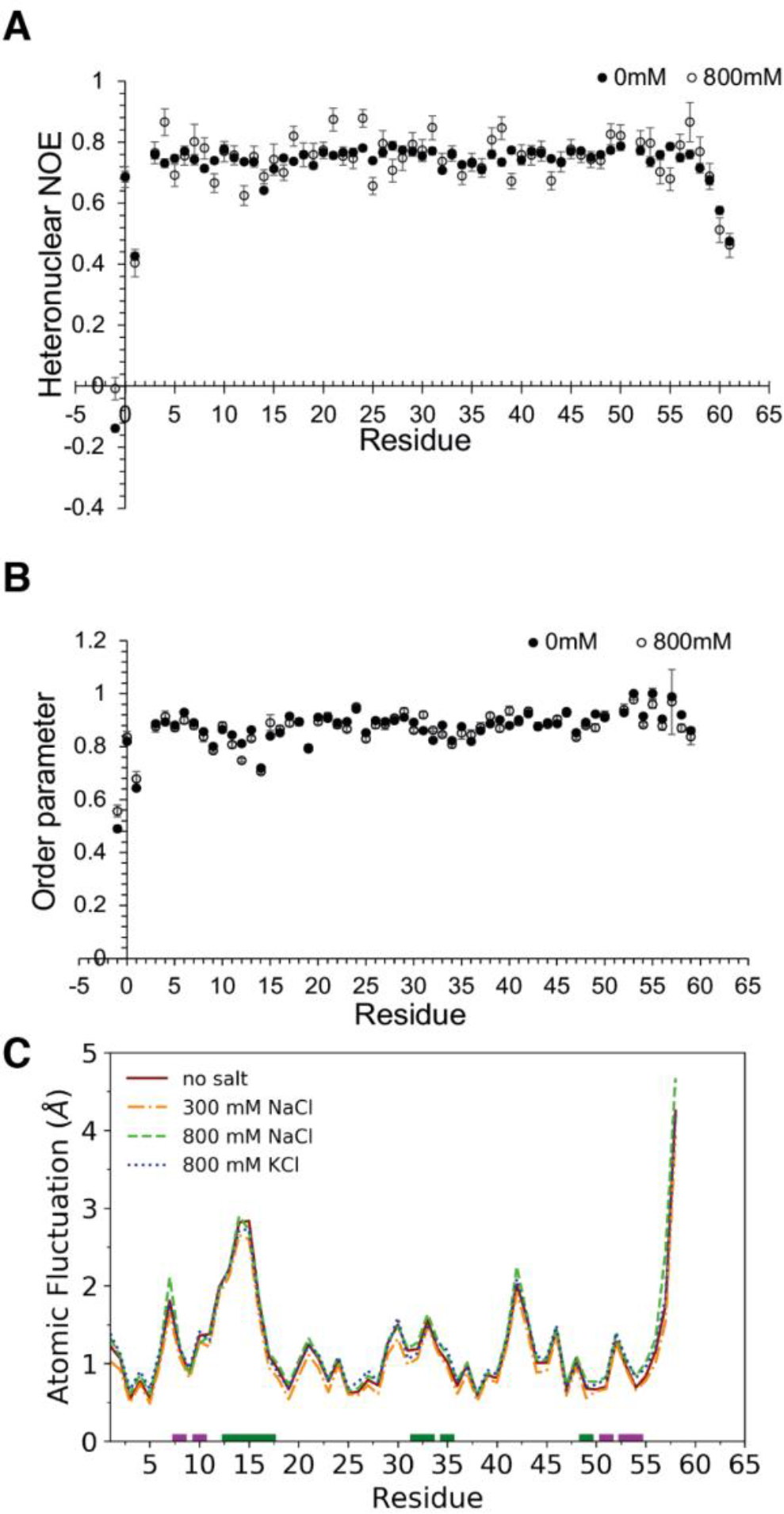
Analysis of domain backbone dynamics. **A**. ^1^H-^15^N heteronuclear NOE of AbpSH3 domain at 0 and 800 mM NaCl. **B**. Backbone ^15^N order parameters calculated from Modelfree. Error bars come from the fit. **C**. Atomic fluctuations of the AbpSH3 domain in no salt, 300 mM NaCl, 800 mM NaCl, and 800 mM KCl simulations. Average standard deviations in the atomic fluctuations (0.2 Å, 0.2 Å, 0.3 Å, and 0.1 Å respectively) are larger than the difference between the lines.

### Intramolecular salt bridges are partially destabilized by salt

MD simulations of the AbpSH3 domain also allow us to measure the intramolecular electrostatic interactions between the negatively charged Asp and Glu residues on the domain and the three positively charged Lys residues. These Lys residues can form intramolecular salt bridges with nearby Asp or Glu residues, which likely stabilize the tertiary structure of the domain. In simulations with 800 mM NaCl and KCl, these intramolecular electrostatic interactions are only slightly disrupted due to the screening of electrostatic interactions by the salt ions (**Fig. 5**).

**Figure 5.**
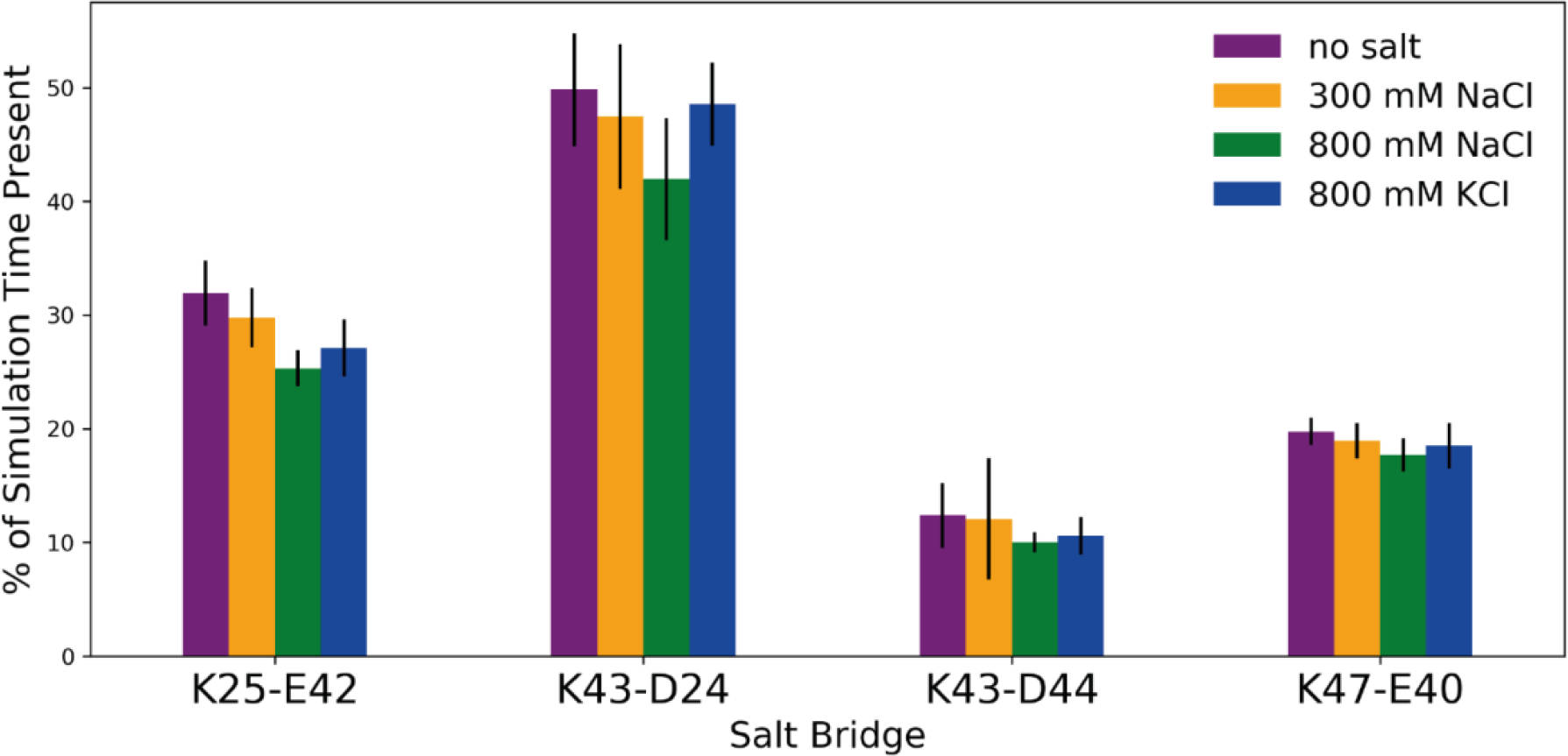
Intramolecular salt bridges. Presence of intramolecular salt bridges in the AbpSH3 domain in simulations with no salt, 300 mM NaCl, 800 mM NaCl, and 800 mM KCl. Salt bridges are considered formed when the two side-chain charged groups are within 4 Å of each other. Error bars represent the standard deviation between the independent simulations.

Consistent with this, MD simulations that measured residue-ion contacts (**Fig 3**) also show that residues involved in intramolecular salt bridges spend less time directly interacting with salt ions. However, these low-frequency interactions with ions (present in 5-30 % of the simulated ensemble) do likely contribute to the small decrease in salt bridge formation observed in the presence of NaCl. Potassium cations disrupt the intramolecular salt bridges less than NaCl (**Fig. 5**), consistent with their less frequent direct binding to the acidic SH3 residues (**Fig 3**).

### Folding rate is very slow due to electrostatic repulsion and is driven by hydrophobic collapse

At low ionic strength, we hypothesized that the conversion of the AbpSH3 denatured state to the transition state (D→‡) would be marginal but must be offset by favorable hydrophobics to compensate for electrostatic repulsion we have measured in the native state. To characterize D→‡, we measured the unfolding kinetics of AbpSH3 at various concentrations of the nonionic chaotrope urea in the absence or presence of NaCl, as well as measuring the unfolding kinetics at 9 M urea at various NaCl concentrations. The data at various urea concentrations were fit to a straight line, representing the unfolding arm of a Chevron plot, allowing us to directly calculate the unfolding rates 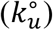 and their kinetic slopes (m_u_) in the absence of urea in 10 mM Tris, pH 8.1, with or without 800 mM NaCl. Folding rates 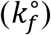 and their kinetic slopes (m_f_) were then determined from this unfolding data and the equilibrium data. The natural log of unfolding rates in 9 M urea were linear with the square root of ionic strength, consistent with the equilibrium data and the Debye-Huckel limiting law (**Fig. 6a**). The effect of salt on the stability of the domain is predominantly due to a 627-fold increase in the folding rate with little change in the unfolding rate (**Table 1**). The increased folding rate in high salt allows us to consider the role of repulsive electrostatic interactions in low salt. All the long-range electrostatic repulsion that destabilizes AbpSH3 at low ionic strength must be present in the transition state, explaining the very slow folding rate of AbpSH3. In comparison, it is in the lowest 10 % of all folding rates recorded for 2-state proteins in the databank of protein folding kinetics (26). In fact, the stability of the transition state improves more than the native state in the presence of salt 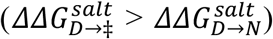, indicating that the net electrostatic repulsion is stronger in the transition state than in the native state, with a 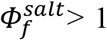. This is consistent with the folding kinetics of a Fyn SH3 domain mutant, where using a 600 mM NaCl change, they measured folding and unfolding rates that produce 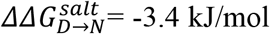 and 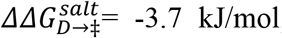, also yielding a 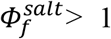 (27). In addition, a study that optimized electrostatic interactions on the surface of the Fyn SH3 domain found that it was stabilized primarily through an eight-fold acceleration in the folding rate (12). Despite the presence of the electrostatic repulsion during AbpSH3 folding 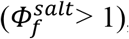, the β_urea_ value without salt is also high at 0.92 (**Table 1**), indicating that indeed D→‡ involves almost complete hydrophobic collapse.

**Table 1.**
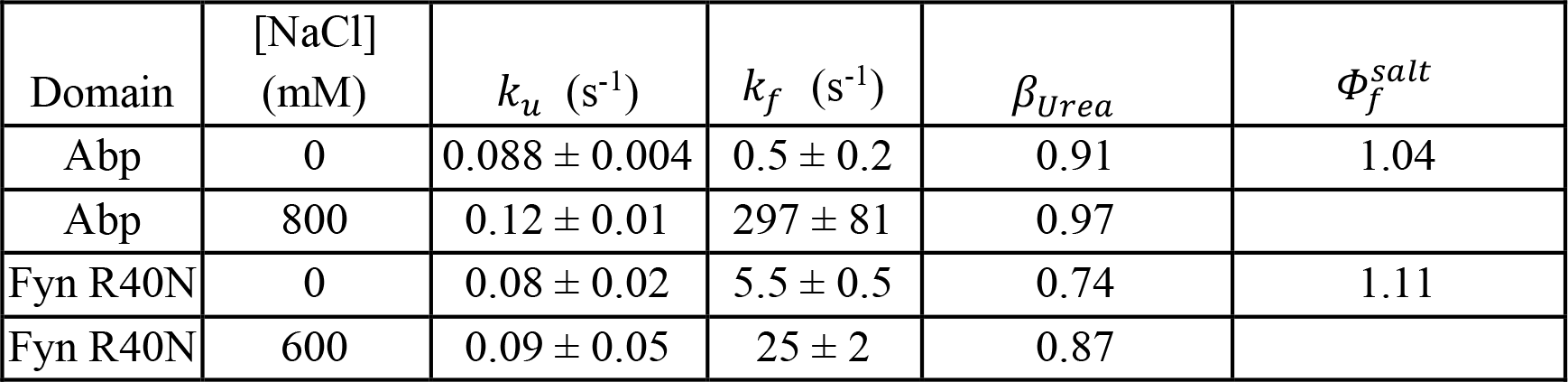
Effect of NaCl on folding kinetics and thermodynamic stability of AbpSH3 and Fyn SH3 domain. Data for Fyn from (27).

| Domain | [NaCl]<br>(mM) | $k_u$ (s<sup>-1</sup>) | $k_f$ (s<sup>-1</sup>) | $\beta_{Urea}$ | $\Phi_f^{salt}$ |
| --- | --- | --- | --- | --- | --- |
| Abp | 0 | $0.088 \pm 0.004$ | $0.5 \pm 0.2$ | 0.91 | 1.04 |
| Abp | 800 | $0.12 \pm 0.01$ | $297 \pm 81$ | 0.97 | |
| Fyn R40N | 0 | $0.08 \pm 0.02$ | $5.5 \pm 0.5$ | 0.74 | 1.11 |
| Fyn R40N | 600 | $0.09 \pm 0.05$ | $25 \pm 2$ | 0.87 | |

**Figure 6.**
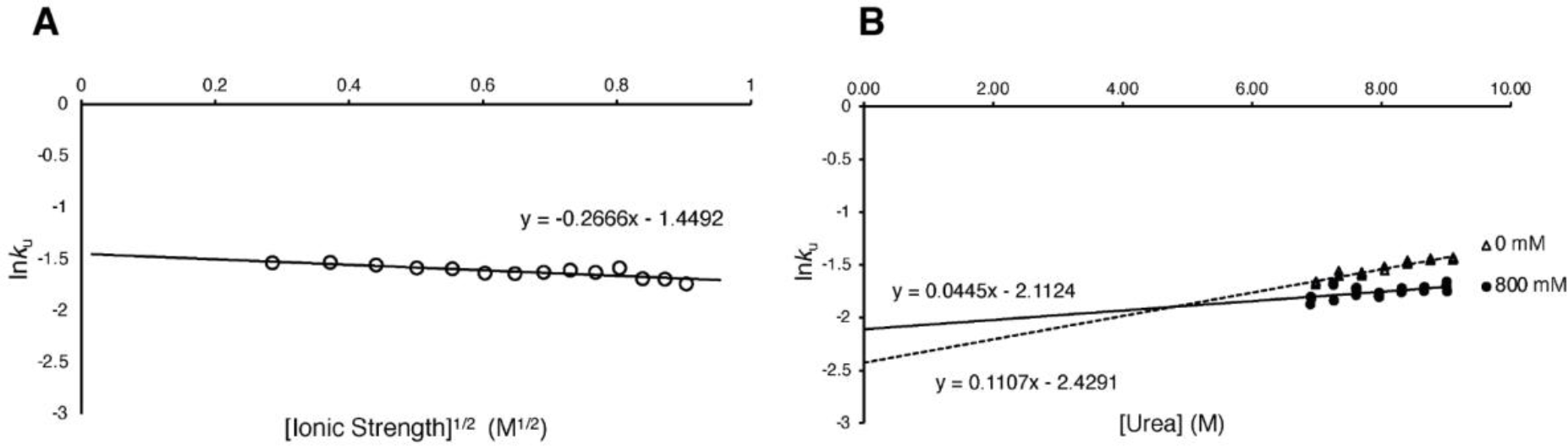
Folding kinetics for AbpSH3 in Urea. **A**. Unfolding rates of AbpSH3 in 9 M urea at various concentrations of NaCl (80 to 800 mM). **B**. Unfolding rates of AbpSH3 at 0 and 800 mM NaCl in various concentrations of urea (7 to 9 M).

### Hydrogen bonds and intramolecular salt bridges form after the transition state

We now consider what drives ‡→N formation. Previous studies (13) on multiple SH3 domains show that many hydrogen bonds between β-strands are first formed during ‡→N as the transition state does not form a complete beta-barrel. Experimentally, it is difficult to probe hydrogen bond formation with a chemical, but we can probe attractive electrostatic interactions using salt. For example, our chemical denaturation data shows that ‡→N at low ionic strength involves some favorable electrostatic interactions, as unfolding is 1.4 times slower compared to high ionic strength. Consistent with this, MD indicates that intramolecular salt bridges are favored in low ionic strength conditions. It is likely these interactions can only occur after transition state formation as the tertiary structure is not fully formed in the transition state. Interestingly, these specific intramolecular salt bridges are found away from the binding site of AbpSH3, not interfering with this negatively charged region, and contributing to a net dipole moment of 242 Debyes across the domain that facilitates fast peptide binding, which has been observed for other SH3 domains (28). It should be noted that as the β_urea_ value without salt is slightly less than 1 (value of 0.92), the conversion of the transition state to the native state (‡→N) at low ionic strength also involves some small additional changes in hydrophobic packing.

Inspecting our chevron unfolding arms (**Fig. 6b)** also allows us to consider unfolding rates at our 2 salt concentrations at high urea concentrations. Our unfolding chevron lines for each salt concentration are not parallel (i.e. different m-values) indicating that 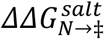that depends on urea concentration, which results from urea’s ability to disrupt attractive electrostatic interactions (for further discussion of this, see supplementary material). Therefore, in addition to the repulsive interactions formed during D→‡ as AbpSH3 folds, additional repulsive electrostatic interactions are formed during ‡→N, which are masked at low urea concentrations because their free energy contribution is smaller than the attractive interactions formed during ‡→N. The same phenomenon was observed with Chicken Fyn SH3 domain with the R40N mutation (27) as reflected by a similar 50 % reduction of the m_u_ value at high ionic strength, revealing differences in 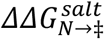that also depend on urea concentration (**Fig. S4**). An earlier study of a truncated FynSH3 protein reported different effects of salt on the unfolding rate, which may be a result of using an N-terminal truncation construct (29). Despite the disruption of attractive electrostatic interactions in 0.8 M salt, unfolding is still quite slow which is likely due to the presence of hydrogen bonds, which are shorter-range interactions that are not affected much by salt (**Table S3**).

We can now consider the driving forces for both steps. Starting at U, hydrophobic collapse drives U→‡ overcoming repulsive electrostatic interactions, while the formation of hydrogen bonds appears to drive ‡→N, with a smaller contribution from intramolecular salt bridges and additional hydrophobic packing (**Fig 7**). This leads to slow unfolding rates at both low and high ionic strength (the rates are similar to the average for 2-state proteins in the databank of protein folding kinetics (26)). NMR and MD data support this, as they show the presence of hydrogen bonds and intramolecular salt bridges in the native state at both low and high ionic strength. Furthermore, there is only a small increase in the β_urea_ value at 800 mM NaCl (0.91 to 0.97), suggesting the transition state structure is similar at different ionic strengths.

**Figure 7.**
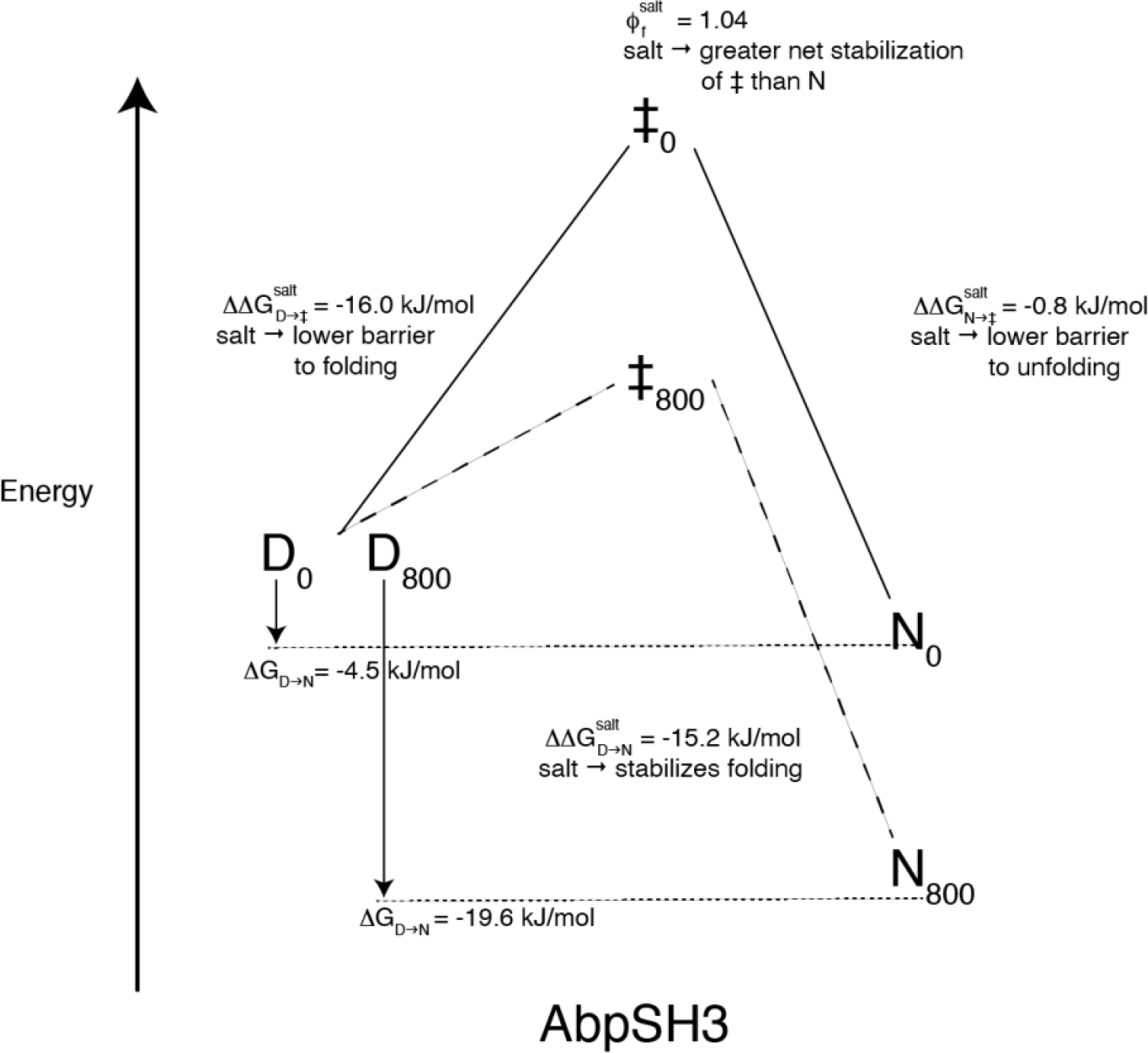
Summary of folding kinetics and thermodynamics for AbpSH3 (SC) at 0 and 800 mM NaCl. The solid line represents 0 NaCl and the dashed line 800 mM NaCl.

Taken together, it is likely that the transition state resembles an electrostatically destabilized dynamic molten-globule with few geometrically stringent interactions due to the flexibility of hydrophobic and long-range electrostatic interactions. This is consistent with a compact, partially-folded, three-stranded sheet for the transition state that has been established in previous studies on other SH3 domains (30-32). It is possible that AbpSH3 has evolved a high degree of hydrophobic collapse in the transition state to compensate for the negative charge repulsion experienced during D→‡. In fact, AbpSH3 has one of the highest β_urea_ values of over ninety 2-state folding proteins (26). This unique collapsed yet electrostatically destabilized transition state is likely to be conserved over evolution as the high net negative charge is present in AbpSH3 orthologs that span over 1 billion years.

## Conclusion

The use of urea and NaCl in equilibrium and kinetic experiments provides complementary information about AbpSH3 folding and which non-covalent forces affect the transition and native states that is further supported by MD and NMR. As hypothesized, we find that at low ionic strength AbpSH3 is folded yet unstable due to electrostatic repulsion and can be stabilized by the addition of salt (**Fig. 7**). This electrostatic repulsion causes slow folding that is driven by hydrophobic collapse, but the repulsion has little effect on the unfolding rate. Instead, hydrogen bonds and intramolecular salt bridges help stabilize the native state, ensuring slow unfolding. Our kinetics data therefore indicate that the transition state resembles a compact molten globule with high electrostatic repulsion compensated somewhat by hydrophobic collapse. Electrostatic repulsion has likely been maintained over evolution to allow fast peptide binding at the cost of slow folding rates (15), which is likely to be the case for almost all SH3 domains. Like other proteins containing conserved residues despite their detrimental effect on domain stability (33), functional advantage has allowed SH3 domains to evolve and maintain an unusually high net charge for such a small protein domain.

## Methods

### Protein expression and purification

A pET32b-based (Novagen) plasmid containing the AbpSH3 domain from from *S*.*cerevisiae* (AbpSC) with a C-terminal his-tag had the following sequence: MAPKENPWATAEYDYDAAEDNELTFVENDKIINIEFVDDDWWLGELEKDGSKGLFPSNYVSLGNLEHHHHHH and the AbpSH3 from the distant orthologue *B*.*dendrobatidis* (AbpBD) had the following sequence: MGYREATAVYDYVAAEPNELSFNEGNLITHIVFVSEDWWQGTLNGVVGLFPGNYVELK QLEHHHHHH. For equilibrium and chemical denaturation assays, the proteins were expressed in autoinduction media and Ni-NTA, followed by purification under denaturing conditions as previously described (34). Proteins were further purified using ion-exchange with a HiTrap-Q column (Cytiva) to >95 % purity and dialysed in 10 mM Tris, pH 8.1 buffer. For ^15^N isotope labeling, AbpSC was expressed using ^15^NH_4_Cl in M9 minimal medium supplemented with trace elements (35), and purified by nickel-affinity chromatography as above. The protein was further purified using Sephadex S200 size-exclusion chromatography (Cytiva) to >95 % purity. For NMR experiments, AbpSC was concentrated to 0.2 - 2 mM and prepared in the following buffer: 1X protease inhibitor, 0.05 % NaN3, 10 % D2O, 10 mM Tris pH 8.1, and different concentrations (0 mM, 100 mM, and 800 mM) of NaCl. For further details see supplementary material

### NMR Spectroscopy

All NMR data was collected using a 600 MHz (14.1T) NMR Agilent DD2 spectrometer equipped with a z-axis gradient HCN room temperature probe at 20 °C. NMR data were processed and analyzed with NMRPipe, ccpnmr analysis, and in-house scripts that made use of the nmrglue python library. Since SH3 domains are small proteins, approximately 60 amino acids, and there are two solution-state NMR structures of AbpSH3, assignments at the various salt concentrations were checked by collecting and analyzing the following spectra H-^15^N HSQC, ^1^H-^15^N TOCSY, and ^1^H-^15^N NOESY. 2D ^1^H-^15^N HSQC spectra were used to calculate the chemical shift differences between the different concentrations of salt using the following equation: Δ_*ppm*_=√((*N*_*ppm*_*shift*∗0.1)^2^+(*H*_*ppm*_*shift*)^2^), where N_ppm_ shift is the difference in ppm in the nitrogen dimension, and H_ppm_ shift is the difference in ppm in the hydrogen dimension. Backbone ^15^NH R_1_, R_1ρ_, and ^1^H-^15^N NOE relaxation data for AbpSC were collected in 10 mM Tris, pH 8.1 with 0 mM, 100 mM, and 800 mM NaCl at 20 °C. ^15^NH R_1_ and R_1ρ_, data were calculated from six timepoints ranging from 1 to 400 ms (R_1_) or 1 to 60 ms (R_1ρ_). R_2_ values were calculated from R_1_ and R_1ρ_ using the relationship R_1ρ_ R_1_ cos^2^θ + R_2_ sin^2^θ, where θ = tan^-1^(ν_SL_/Ω), ν_SL_ is the strength of the applied ^15^NH spin lock (∼2000 Hz), and Ω is the offset of the peak from the ^15^N carrier. Errors in the relaxation rates were derived from the covariance matrix of the fit. Backbone ^15^NH R_1_, R_2_, and NOE data were used to calculate residue specific order parameters (S^2^) and an effective correlation time for internal motion (τ_e_) as well as the overall global rotational correlation time (τ_c_) using the model-free approach as implemented in Modelfree v4.2 (https://comdnmr.nysbc.org/comd-nmr-dissem/comd-nmr-software/software/modelfree). All data were fit using “model 2” using an axially symmetric diffusion model resulting in an average τ_c_ value of approximately 5.76 +/- 0.03 ns across the salt concentrations. N-H bond lengths of 1.02 Å and ^15^N chemical shift anisotropy of -160 ppm were used in the fitting. All assignment data at 0, 100 and 800 mM NaCl have been deposited in BMRB with IDs 51744, 51745, 51746.

### Equilibrium and kinetic chemical denaturation measurements

All experiments were performed in a POLARstar Omega plate reader (BMG labtech, NC, USA) with a Hellma 96 well quartz plate at 30 °C. The assays were performed as previously described (36) although modified to avoid injecting denaturants into the wells. For the equilibrium assay, in each well, 50 µL of protein started in high concentrations of urea (between 6 and 10.5 M) in 10 mM Tris, pH 8.1 (+ salt) and were refolded by injection (dilution) using the same buffer, without urea. Fewer samples were measured at a time to enable a complete set of data for a given protein to take around 2 hours, minimizing the effects of evaporation from the wells. For any given urea concentration, multiple time points were collected and analyzed to ensure equilibrium had been reached before the next urea injection was made. Errors from the equilibrium assay were calculated as standard deviations from at least 3 separate measurements. Errors of slopes and intercepts were calculated from least-squared regression analysis. For the kinetic assay, 10 µL of 50 µM of protein in 10 mM Tris, pH 8.1 (+ salt) was injected into wells with 90 µL of 10 mM Tris, pH 8.1 (+ salt) with different concentrations of urea. Unfolding was monitored by tryptophan fluorescence for 1 minute per well to also minimize the effects of evaporation from the well. k_u_ was determined from the unfolding arm of the chevron plot while k_f_ was determined from k_u_ and K_eq_ as described previously. Errors for k_u_ and m_u_ were calculated by least-squared regression analysis and errors for k_f_ by error propagation. For the details of the thermodynamic and kinetic equations used, see supplementary material.

### Molecular Dynamics Simulations

#### Simulations

To study the electrostatic interactions in the AbpSH3 complex with atomic resolution, molecular dynamics (MD) simulations of the domain were run under different salt conditions. All simulations were initiated from the AbpSH3 crystal structure (PDB: 1JO8) (37) and were run under conditions of 0 mM salt, 300 mM sodium chloride, 800 mM sodium chloride, and 800 mM potassium chloride. Simulations were run on a local cluster with Amber 16 using the Amber ff14SB force field and the CUDA version of pmemd for GPUs (38, 39). All systems were solvated with TIP3P-FB water (40). When solvated, all structures were placed at least 9 Å from the edges of the water box.

For the simulations with 300 or 800 mM salt, the number of cations needed to reach the appropriate concentration was first calculated based on the volume of the water box before equilibration. Using the *LEaP* module within the AmberTools16 package (38), the calculated number of cations were added as well as the appropriate number of chloride anions needed to neutralize the overall charge of the system. **Table S4** indicates the number of ions added to each simulation. Low salt simulations only included enough sodium ions to neutralize the system.

Before running production, all systems were subjected to minimization, heating, and equilibration. Two rounds of minimizations were completed without restraints at 1000 steps each. Each round consisted of 500 cycles of steepest descent minimization and 500 cycles of conjugate gradient minimization. The temperatures of all systems were then raised from 100 K to 300 K over 40 ps of simulation time with a 10 kcal/mol harmonic potential force constant restraint on the protein atoms. All systems were then equilibrated twice in the NPT ensemble at 1.013 bar, first for 50 ps with the same restraints on the protein and the Berendsen barostat and again for 200 ps without restraints using a Monte Carlo barostat. Bonds to hydrogen were constrained using the SHAKE algorithm. The particle-mesh Ewald procedure was used to handle long-range electrostatic interactions with a non-bonded cutoff of 9 Å for the direct space sum. Independent production simulations were then run using random starting velocities for 1.0-2.7 μs each. Production simulations were run in the NPT ensemble at 300 K and 1.013 bar, using Langevin dynamics with a collision frequency of 1.0 ps^-1^, with a Monte Carlo barostat, new system volumes attempted every 100 steps, an integration step every 2 fs, and coordinates stored every 10 ps. **Table S4** indicates the number of independent simulation runs for each system and the total simulation time for each system.

### Analysis

The *cpptraj* module in AmberTools16 along with python scripts created by our group were used to conduct atom-to-atom distance measurements, atomic fluctuations analysis, and hydrogen bond analysis (38). For ion contacts and salt bridges, the distance between the positively charged nitrogen atom on the lysine sidechain and the most distal carbon atom (bonded to a negatively charged oxygen atom) of glutamate or aspartate was measured. These distance measurements were used to calculate the percent of time that certain salt bridges and contacts with salt ions were present in the simulations. A cutoff distance of 4 Å was used for salt bridges and salt ion contacts with charged residues. The average values from each independent simulation were used to calculate standard deviation values which are shown as error bars on the plots. Structure figures were created with Visual Molecular Dynamics (41).

## Supplementary Material

File name: Supplementary_Material.pdf, PDF document, 621 MB

Supplementary Method. Minimal Media Expression.

Supplementary Method. Thermodynamics and kinetics of folding.

Supplementary Discussion. Effect of urea on electrostatic interactions.

Figure S1. Free energy of folding vs ionic strength shows a weaker fit to lines.

Figure S2. Free energy of folding vs log [ionic strength] shows a weaker fit to lines.

Figure S3. Salt has an effect in and around most charged residues on the AbpSH3 domain.

Figure S4. Full Chevron plots at two NaCl concentrations for AbpSC and Fyn R40N.

Table S1. Ionic strength relationships with stability for AbpSH3 from *S*.*cerevisiae* (SC) and *B*.*dendrobatidis* (BD).

Table S2. Data summary of kinetic and equilibrium folding data for AbpSH3 (SC) and FynSH3 R40N.

Table S3. Average number of intramolecular hydrogen bonds formed in simulations.

Table S4. Simulation data for each system analyzed.

File name: FileS1.xlsx, Excel file, 128 KB

Supplementary Data. Covariation data across AbpSH3 orthologs.

## Author Contributions

EJS conceived the idea for this research. EJS, VJ, GMB, MES, MPL, MD, BL collected experimental data. AHC, MAC, and AJA collected simulation data. EJS, VJ, MD analyzed experimental data. AHC, MAC, and KAB analyzed simulation data. EJS and KAB wrote the paper.

## Acknowledgements

The authors thank Kristina Foley, Colin McClure and Zoey Sharp for important preliminary data collection and analysis. Thank you to Michael Donnelly for computational support. KAB thanks the MERCURY Consortium for mentoring support. Research reported in this publication was supported by an Institutional Development Award (IDeA) from the National Institute of General Medical Sciences of the National Institutes of Health under grant number P20GM103451 (EJS), a Wellcome Trust Summer Internship (MES), the National Science Foundation award MCB-1852677 (KAB) and NIH grant R35GM128906 (MPL). AHC was supported by the Skidmore Schupf Scholar Program.

