## Supplemental Material for "How a highly acidic SH3 domain folds in the absence of its charged peptide target"

### Supplementary Material for How a highly acidic SH3 domain folds in the absence of its charged peptide target

#### Minimal Media Expression

For minimal media expression, AbpSC was separately transformed into *Escherichia coli* BL21 gold cells. A single colony was used to inoculate 30 mL of TB growth media containing 100 μg/mL ampicillin and shaken for 5 hours. Cells were spun at 1800 rpm for 10 minutes and resuspended in 100 mL of $^{15}NH_4Cl$ in M9 minimal medium and grown overnight. The cells were spun at 1800 rpm for 10 minutes and resuspended in $^{15}NH_4Cl$ in M9 minimal medium supplemented with trace elements (1), shaking at 37 °C until an $OD_{600}$ of 1. Cells were then induced with isopropyl-β-D-1-thiogalactopyranoside (IPTG) at a final concentration of 1 mM and shaken at 37 °C until $OD_{600}$ was close to 2. Next, the cells were spun at 4600 rpm for 20 minutes. The pellet was resuspended in 15 mL of leftover minimal media and spun again at 10,000 rpm for 2 minutes. The pellet was resuspended in lysis buffer (20 mM Tris, 300 mM NaCl, 10 mM imidazole, pH 8.0) with 0.4 mg/mL lysozyme, 1X protease inhibitor, 0.1 mg/mL DNase and 1 % Triton X-100. The cell suspension was incubated at 4 °C for 45 minutes, followed by centrifugation for 20 minutes at 12000 rpm to remove the insoluble material. Protein purification was performed by nickel-affinity chromatography using a multi-column plate adapter as previously described (2). The protein was further purified using Sephadex S200 size-exclusion chromatography (Cytiva) to >95 % purity. For NMR experiments, AbpSC was concentrated to 0.2 - 2 mM and prepared in the following buffer: 1X protease inhibitor, 0.05 % NaN3, 10 % D2O, 10 mM Tris pH 8.1, and different concentrations (0 mM, 100 mM, and 800 mM) of NaCl.

#### Thermodynamics and kinetics of folding

The free energy change for protein folding in the absence of denaturant in 10 mM Tris, pH 8.1 at 30 °C is defined as $\Delta G^{\circ}_{D\rightarrow N}$. This is related to the protein folding equilibrium constant, $K$, which is the ratio of the protein folding rate constant $k_f$ and the protein unfolding rate constant $k_u$ according to

**Equation 2:**

$$\Delta G^{\circ}_{D\rightarrow N} = -RTln(K) = -RTln\left(\frac{k_f}{k_u}\right) = -RT\left[ln\left(k_f\right) - ln\left(k_u\right)\right] .$$

Denaturant is used in our assays to directly measure the equilibrium constant, *K*, and rate constant value for unfolding $k_u$. Using equation 2 with these measured values, we can then calculate the rate constant for folding, $k_f$.

The free energy change for protein folding at any given urea concentration, $\Delta G^{urea}_{D\rightarrow N}$ is linearly dependent on urea concentration with a slope of $m_{eq}$ according to

**Equation 3a:**

$$\Delta G^{urea}_{D\rightarrow N} = \Delta G^{\circ}_{D\rightarrow N} + m_{eq}[Urea]$$

where the y-intercept of the line, $\Delta G^{\circ}_{D\rightarrow N}$, described in this equation represents the standard free energy change for protein folding in the absence of urea.

At the [Urea] where 50% of the protein is unfolded (the midpoint in an equilibrium unfolding curve), $D_{50}$, $\Delta G^{urea}_{D\rightarrow N}$ = 0 and therefore

**Equation 3b:**

$$\Delta G^{\circ}_{D\rightarrow N} = -m_{eq}D_{50}$$

Because urea stabilizes solvent exposed hydrophobic surfaces in a denatured protein, the magnitude of the $m_{eq}$ value is correlated with the extent that the protein solvent accessible surface area changes upon unfolding. For more stable proteins, where the folded state is more collapsed compared to the denatured state, the $m_{eq}$ value is larger and therefore urea has a greater ability to destabilize the protein. Both $D_{50}$ and $m_{eq}$ are determined from the fit to equilibrium data from our equilibrium assay (3) allowing us to calculate $\Delta G^{\circ}_{D\rightarrow N}$ directly.

Protein folding kinetic measurements directly yield observed folding or unfolding rate constants, $k_f$ and $k_u$, at different final urea concentrations. The plots of $\ln(k_f)$ and $ln(k_u)$ versus urea concentration are linear, with different slopes, $-m_f$ and $m_u$ , respectively as shown in the equations

**Equation 4a:**

$$ln(k_u^{urea}) = ln(k_u^{\circ}) + m_u[Urea]$$

**Equation 4b:**

$$ln\left(k_f^{urea}\right) = ln\left(k_f^{\circ}\right) - m_f[Urea].$$

where $k_f^{\circ}$ and $k_u^{\circ}$ are folding rate constants in the absence of urea. These can be rearranged to give expressions for $m_f$ and $m_u$:

**Equation 4c:**

$$ln\left(\frac{k_u^{urea}}{k_u^{\circ}}\right) = m_u[Urea] \text{ and}$$

**Equation 4d:**

$$ln\left(\frac{k_f^{urea}}{k_f^{\circ}}\right) = -m_f[Urea].$$

Sometimes only one of these rate constants is convenient to measure directly, but by using the equilibrium relationship in equation 2, either the folding or unfolding rate constant can be used to calculate the other rate constant. It is also useful to define the relationship between $m_{eq}$, $m_u$and $m_f$. Rearranging Equation 3 we can define $m_{eq}$ in terms of folding and unfolding rates.

**Equation 5:**

$$m_{eq}[Urea] = \Delta G_{D\rightarrow N}^{urea} - \Delta G_{D\rightarrow N}^{\circ}$$

and, using equation 2,

$$\Delta G_{D\rightarrow N}^{urea} - \Delta G_{D\rightarrow N}^{\circ} = -RTln(K^{urea}) - [-RTln(K^{\circ})]$$

$$= -RTln\left(\frac{K^{urea}}{K^{\circ}}\right) = RTln\left(\frac{k_u^{urea}}{k_u^{\circ}}\right) - RTln\left(\frac{k_f^{urea}}{k_f^{\circ}}\right).$$

Now we can substitute equations 4c and 4d into equation 5 to generate an equation for $m_{eq}$ in terms of $m_u$ and $m_f$ yielding,

$$m_{eq}[Urea] = RTm_u[Urea] + RTm_f[Urea]$$

and therefore

**Equation 6:**

$m_{eq} = RT(m_u + m_f)$.

Equation 6 allows us to calculate $m_f$ using $m_u$ and $m_{eq}$ which are measured directly.

Because $m_f$ and $m_{eq}$ measure the effect of urea on the folding rate and free energy of folding respectively, the ratio of these values captures the effect of urea on the transition state formation compared to the formation of the folded state. Thus, the degree to which the protein is collapsed in the transition state compared to the folded state can be quantified by this ratio through calculation of the beta tanford value, $\beta_{Urea}$, using equation 7.

**Equation 7:**

$\beta_{Urea} = \frac{RTm_f}{m_{eq}}$.

This equation can be derived by starting with

**Equation 8a:**

$$\beta_{Urea} = \frac{\Delta\Delta G^{Urea}_{D\rightarrow\ddagger}}{\Delta\Delta G^{Urea}_{D\rightarrow N}},$$

where $\Delta\Delta G^{Urea}_{D\rightarrow N}$, is the difference in the free energy change for $\Delta G_{D\rightarrow N}$ in the presence and absence of a given urea concentration, and $\Delta\Delta G^{Urea}_{D\rightarrow\ddagger}$ is the difference in free energy change to reach the transition state from the denatured state for that same urea concentration. Since the difference in free energy between the denatured state, *D*, and the native state, *N*, is equal to the free energy to go from *D* to the transition state, ‡, plus the free energy to go from ‡ to *N*, we can also write

**Equation 9:**

$$\Delta\Delta G^{Urea}_{D\rightarrow N} = \Delta\Delta G^{Urea}_{D\rightarrow\ddagger} + \Delta\Delta G^{Urea}_{\ddagger\rightarrow N} .$$

Similar to the free energy change for protein folding, $\Delta G^{\circ}_{D\rightarrow N}$, the free energy change from *D* to ‡ can be expressed in terms of rate constants for folding, $k_f$, and for the decay of the transition state, $k_{\ddagger}$, as seen in equation 2, according to

**Equation 10:**

$$\Delta G^{\circ}_{D\rightarrow\ddagger} = -RTln(K_{\ddagger}\ ) = -RTln\left(\frac{k_f}{k_{\ddagger}}\right) = -RT\left[ln(k_f\ ) - ln(k_{\ddagger}\ )\right],$$

and the expression for $\Delta G^{\circ}_{N\rightarrow\ddagger}$ can be expressed similarly in terms of $k_u$ and $k_{\ddagger}$. The difference in the free energy change for $\Delta G^{\circ}_{D\rightarrow\ddagger}$ in the presence and absence of urea, $\Delta\Delta G^{urea}_{D\rightarrow\ddagger}$, is therefore given by

**Equation 11a:**

$$\Delta\Delta G^{urea}_{D\rightarrow\ddagger} = -RT\left[ln(k^{urea}_f) - ln(k^{urea}_{\ddagger})\right] + RT\left[ln(k^{\circ}_f) - ln(k^{\circ}_{\ddagger})\right].$$

By definition, the rate of decay of transition state, $k_{\ddagger}$, is equal in both directions. If we assume $k_{\ddagger}$ does not depend on the urea concentration, then $k^{urea}_{\ddagger}$ and $k^{\circ}_{\ddagger}$ are equal, and we can write

**Equation 11b:**

$$\Delta\Delta G_{D\rightarrow\ddagger}^{urea} = -RTln\left(k_f^{urea}\right) + RTln\left(k_f^{\circ}\right).$$

Substituting equation 11b and equations 5 and 4b into Equation 8a, we find

**Equation 8b:**

$$\beta_{Urea} = \frac{-RT\left[ln\left(k_f^{Urea}\right) - ln\left(k_f^{\circ}\right)\right]}{\left(\Delta G_{D\rightarrow N}^{Urea} - \Delta G_{D\rightarrow N}^{\circ}\right)} = \frac{RTm_f[Urea]}{m_{eq}[Urea]} = \frac{RTm_f}{m_{eq}}.$$

Substituting equation 6 in Equation 8b, we can also directly calculate $\beta_{Urea}$ using $m_{eq}$ and $m_u$ according to

**Equation 8c:**

$$\beta_{Urea} = \frac{m_{eq} - RTm_u}{m_{eq}}.$$

Thus, by measuring $m_{eq}$ and $m_u$, we can arrive at the beta-tanford value that quantifies the effect of urea on the folding transition state compared to the effect on the overall stability.

#### Effect of salt on folding

In order to quantify the effect of salt on the folding transition state, we can also define a folding phi-value for salt analogous to $\beta_{Urea}$,

**Equation 12a:**

$$\Phi_f^{salt} = \frac{\Delta\Delta G_{D\rightarrow\ddagger}^{salt}}{\Delta\Delta G_{D\rightarrow N}^{salt}}.$$

We can calculate the folding phi-value for salt directly from the folding rates constants in the presence and absence of salt, according to

**Equation 12b:**

$$\Phi_f^{salt} = \frac{-RTln\left(\frac{k_f^{salt}}{k_f^{\circ}}\right)}{\Delta\Delta G_{D\rightarrow N}^{salt}},$$

or by using m-values from our plots as we did for $\beta_{Urea}$.

The free energy of folding in the presence of salt depends linearly on the square root of the ionic strength, as described in the Debye-Huckel equation (equation 1),

**Debye-Huckel:**

$$\Delta G_{N\to D}^{salt} = \Delta G_{N\to D}^{\circ} + m_{eq}^{salt} I^{\frac{1}{2}} \ .$$

The Debye-Huckel equation is analogous to equation 3 and gives us $m_{eq}^{salt}$ for salt from the slope of the plot. Likewise, analogous to equation 4a, we define $m_u^{salt}$ as the slope of the plot of $ln(k_u)$ versus the square root of ionic strength according to

**Equation 13:**

$$ln(k_u^{salt}) = ln(k_u^{\circ}) + m_u^{salt} I^{\frac{1}{2}} \ ,$$

where $k_u^{\circ}$ is the unfolding rate constant in the absence of salt. Using these salt m-values, we can then calculate the phi-value for salt directly:

**Equation 12c:**

$$\Phi_f^{salt} = \frac{m_{eq}^{salt} - RT m_u^{salt}}{m_{eq}^{salt}}.$$

This salt phi-value is also analogous to the classic protein folding mutation phi value. The mutation phi value captures the effect on the transition state compared to the effect on the overall stability when a mutation has been introduced, while the salt phi-value, $\Phi_f^{salt}$, captures the effect on the transition state compared to the effect on the overall stability when salt has been introduced.

#### Effect of urea on electrostatic interactions

At 0 M urea, the $\Delta\Delta G_{N\rightarrow\ddagger}^{salt}$ is -0.8 kJ/mol (difference of y-intercepts) which corresponds to unfolding in low salt being 1.4 times slower compared to high ionic strength since the intramolecular bonds are stronger as discussed above. Surprisingly however, at 9 M urea $\Delta\Delta G_{N\rightarrow\ddagger}^{salt}$ is 0.8 kJ/mol, indicating that in low salt the protein actually unfolds faster than in high salt. In low ionic conditions, increasing urea concentration appears to linearly reduce the attractive electrostatic contribution from intramolecular salt bridges favorable to folding (**Fig. 6b)**. We hypothesize that during the 2-second mixing time of the experiment, urea interacts with the charged residues on the protein surface, disrupting intramolecular salt bridges, before unfolding can take place. In this case, we envision urea interacting with the amino group of the surface lysines through hydrogen bonds before it interacts with the backbone to induce unfolding, consistent with proposed urea interaction mechanisms (4, 5). Therefore, at high urea concentrations where favorable salt bridges are essentially screened by urea interacting with lysines, unfolding is faster at lower ionic strength because repulsive interactions can still exert an effect compared to higher ionic strengths.

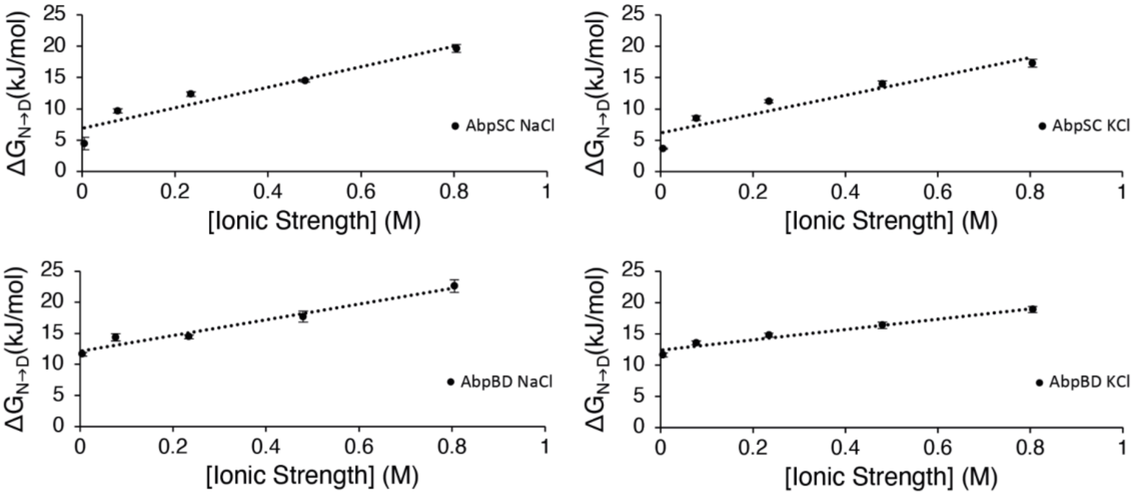


Figure S1. Free energy of folding vs ionic strength shows a weaker fit to lines

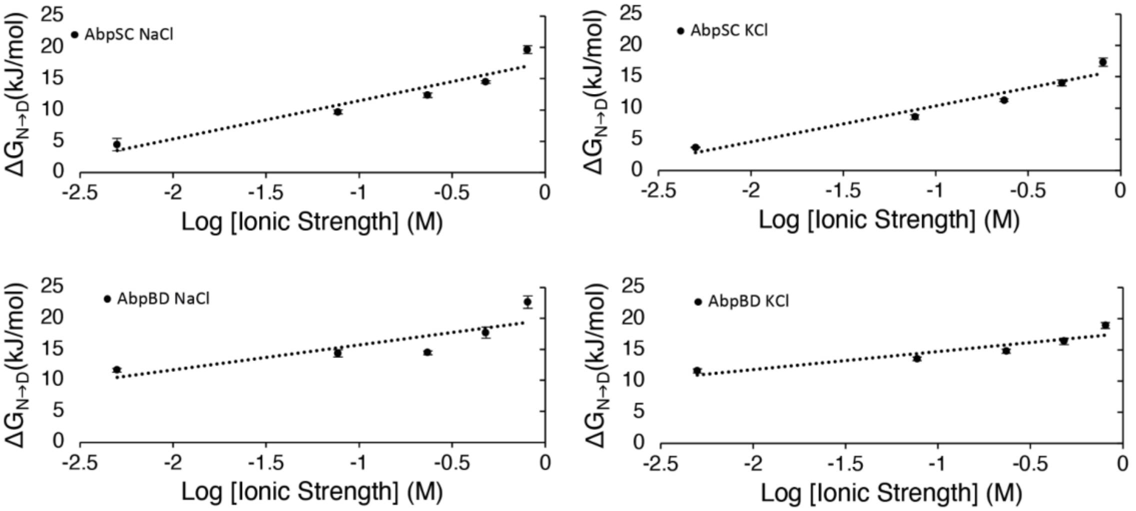


Figure S2. Free energy of folding vs log [ionic strength] shows a weaker fit to lines

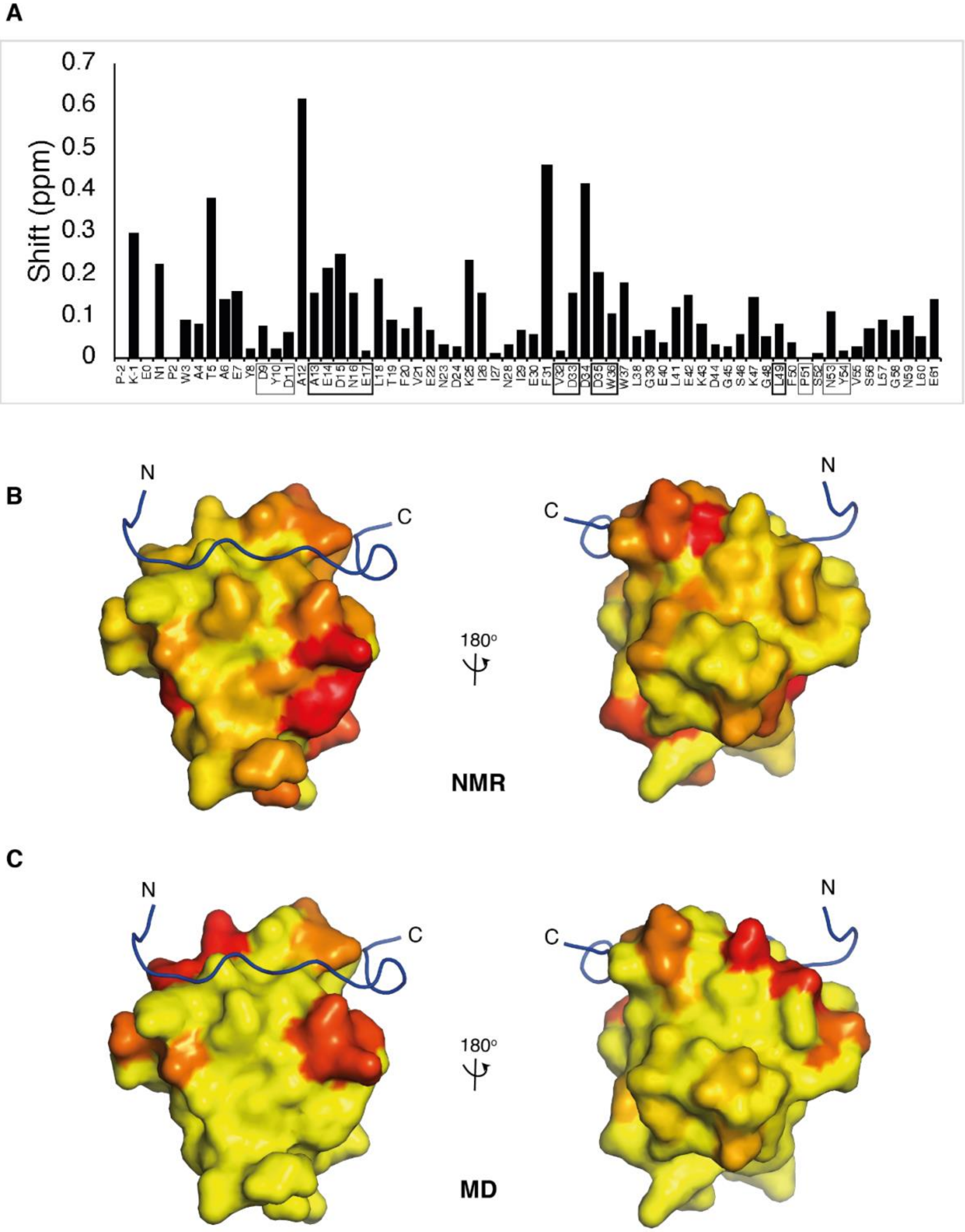
A
0.7
0.6
0.5
0.4
0.3
0.2
0.1
0
Shift (ppm)
P-2 K-1 E0 N1 P2 W3 A4 T5 A6 E7 Y8 D9 Y10 D11 A12 A13 E14 D15 N16 E17 L18 T19 F20 V21 E22 N23 D24 K25 I26 I27 N28 I29 E30 F31 V32 D33 D34 D35 W36 W37 L38 G39 E40 L41 E42 K43 D44 G45 S46 K47 G48 L49 F50 P51 S52 N53 Y54 V55 S56 L57 G58 N59 L60 E61
B
N
C
C
N
180°
NMR
C
N
C
C
N
180°
MD

Figure S3. Salt has an effect in and around most charged residues on the AbpSH3 domain.

***A.*** *Chemical shift differences between 0 and 800 mM NaCl for AbpSH3 mapped onto AbpSH3-peptide complex (2rpn). Residues affected by salt are coloured yellow (no change) to red (0.4 ppm shift or greater).* ***B.*** *Chemical shift differences between 0 and 800 mM NaCl for AbpSH3. Surface I and surface II binding residues are boxed in thin and thick lines, respectively. Overall, from NMR chemical shift changes, salt appears to have an effect on the whole domain. There is a weak overall correlation between the residues shifted by peptide binding and those shifted by high salt. Residues (-2), 2, and 51 are prolines and have no value in this plot.* ***C.*** *MD simulation data used in Figure 3 shown as surface representation where the location of charged residues that interact with ions are displayed on the AbpSH3 NMR structure (2RPN). The ArkA peptide is shown to indicate where it binds, although it was not present in the simulations. The colour scheme is red to yellow to indicate the percent of simulation time that residue is in contact with an oppositely charged ion in the 800 mM NaCl simulations.*

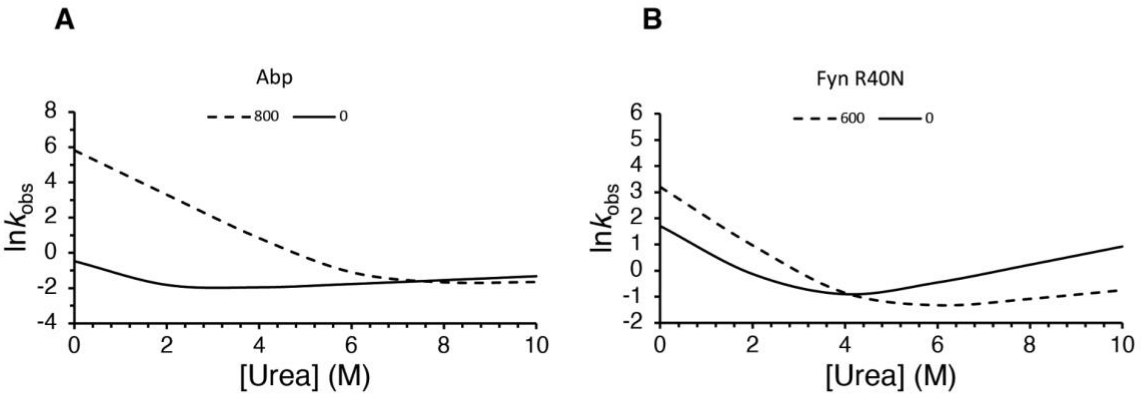


Figure S4. Full Chevron plots at two NaCl concentrations for AbpSC and Fyn R40N.

***A.*** *Chevron plot of AbpSH3 at 0 (solid) and 800 (dashed) mM NaCl, shows salt has a bigger effect on protein folding than unfolding.* ***B.*** *Chevron plot of Fyn R40N at 0 (solid) and 600 (dashed) mM NaCl, shows salt has an equal effect on protein folding and unfolding.*

Table S1. Ionic strength relationships with stability for AbpSH3 from *S.cerevisiae* (SC) and *B.dendrobatidis* (BD).

| Square root of Ionic Strength Plot | | | | |
|---|---|---|---|---|
| Protein | Salt | $m_{eq}^{salt}$ (kJ mol$^{-1}$ M$^{-1}$) | $\Delta G^{\circ}_{D\rightarrow N}$ (kJ mol$^{-1}$) at 0 Salt | $R^2$ |
| AbpSC | NaCl | 17.00 ± 1.60 | 3.90 ± 0.90 | 0.97 |
| AbpSC | KCl | 15.84 ± 1.06 | 3.30 ± 0.60 | 0.99 |
| AbpBD | NaCl | 12.24 ± 2.30 | 10.22 ± 1.30 | 0.90 |
| AbpBD | KCl | 8.34 ± 0.60 | 10.99 ± 0.34 | 0.98 |
| Ionic Strength Plot | | | | |
| Protein | Salt | $m_{eq}^{salt}$ (kJ mol$^{-1}$ M$^{-1}$) | $\Delta G^{\circ}_{D\rightarrow N}$ (kJ mol$^{-1}$) at 0 Salt | $R^2$ |
| AbpSC | NaCl | 16.45 ± 3.00 | 6.87 ± 1.30 | 0.91 |
| AbpSC | KCl | 15.08 ± 3.03 | 6.14 ± 1.31 | 0.89 |
| AbpBD | NaCl | 12.64 ± 1.44 | 12.10 ± 0.62 | 0.96 |
| AbpBD | KCl | 8.25 ± 0.95 | 12.39 ± 0.41 | 0.96 |
| Log of Ionic Strength Plots | | | | |
| Protein | Salt | $m_{eq}^{salt}$ (kJ mol$^{-1}$ M$^{-1}$) | $\Delta G^{\circ}_{D\rightarrow N}$ (kJ mol$^{-1}$) at 0 Salt | $R^2$ |
| AbpSC | NaCl | 6.11 ± 1.16 | 17.59 ± 1.37 | 0.90 |
| AbpSC | KCl | 5.77 ± 0.85 | 16.11 ± 1.01 | 0.94 |
| AbpBD | NaCl | 4.03 ± 1.51 | 19.74 ± 1.79 | 0.70 |
| AbpBD | KCl | 2.89 ± 0.71 | 17.61 ± 0.84 | 0.85 |

Table S2. Data summary of kinetic and equilibrium folding data for AbpSH3 (SC) and FynSH3 R40N.

| | [NaCl] (mM) | $k_u$ (s$^{-1}$) | $k_f$ (s$^{-1}$) | $\Delta G_{D\rightarrow N}$ (kJ mol$^{-1}$) | $\Delta\Delta G_{D\rightarrow N}^{salt}$ (kJ mol$^{-1}$) | $\Delta\Delta G_{D\rightarrow ‡}^{salt}$ (kJ mol$^{-1}$) | $\Delta\Delta G_{N\rightarrow ‡}^{salt}$ (kJ mol$^{-1}$) | $m_u$ (M$^{-1}$) | $m_f$ (M$^{-1}$) | $m_{eq}$ (kJ mol$^{-1}$ M$^{-1}$) | $\beta_{Urea}$ | $\Phi_f^{salt}$ |
|---|---|---|---|---|---|---|---|---|---|---|---|---|
| Abp | 0 | 0.088 ± 0.004 | 0.5 ± 0.2 | -4.5 ± 1 | -15.2 | -16 | -0.8 | 0.11 ± 0.006 | 1.13 ± 0.01 | 3.14 ± 0.03 | 0.91 | 1.04 |
| Abp | 800 | 0.12 ± 0.01 | 297 ± 81 | -20 ± 0.6 | | | | 0.04 ± 0.01 | 1.24 ± 0.05 | 3.2 ± 0.1 | 0.97 | |
| Fyn R40N | 0 | 0.08 ± 0.02 | 5.5 ± 0.5 | -11 ± 0.7 | -3.4 | -3.8 | -0.4 | 0.35 ± 0.04 | 1.01 ± 0.06 | 3.37 ± 0.03 | 0.74 | 1.11 |
| Fyn R40N | 600 | 0.09 ± 0.05 | 25 ± 2 | -14 ± 1 | | | | 0.17 ± 0.08 | 1.14 ± 0.04 | 3.25 ± 0.04 | 0.87 | |

Table S3. Average number of intramolecular hydrogen bonds formed in simulations.

| AbpSH3 | AbpSH3 300mM NaCl | AbpSH3 800 mM NaCl | AbpSH3 300mM NaCl |
|---|---|---|---|
| 26.9 ± 0.5 | 27.2 ± 0.4 | 26.0 ± 0.3 | 27.1 ± 0.4 |

*Uncertainty is based on the standard deviation based on the independent simulations.*

Table S4. Simulation data for each system analyzed.

| System | # of independent simulations | Total simulation time (µs) | # of cations | # of anions | Volume of water box ($nm^3$) |
|---|---|---|---|---|---|
| **AbpSH3 in 0 mM salt** | 10 | 24 | 12 (sodium ) | None | 88 |
| **AbpSH3 in 300mM sodium chloride** | 10 | 10 | 19 | 7 | 87 |
| **AbpSH3 in 800mM sodium chloride** | 10 | 24 | 51 | 39 | 86 |
| **AbpSH3 in 800mM potassium chloride** | 10 | 30 | 51 | 39 | 87 |

File S1. Covariation data across AbpSH3 orthologs.

***Tab 1.*** *262 orthologous fungal Abp1p sequences were obtained from (6).* ***Tab 2.*** *These sequences were analyzed by the covariation analysis program Gremlin at http://openseq.org/submit.php and 6 pairs of residues (24&41, 28&40, 21&24, 6&20, 7&56, 40&47) with a high score (greater than 2), a high probability (greater than 0.9) and in close contact (within 4 angstroms according to pdb structure 2rpn) were found. These residues were color coded according to 6 equivalency groups defined in (6) to look for trends over the sequences. The only salt bridge in the yeast AbpSH3 that has covariation is between residues 40 and 47, however, this charged interaction is not conserved and is frequently between polar interactions. In the other covaried pairs, charge-charge interactions are not conserved either.*
